# Upfront Menin-inhibitor resistance in multiply pretreated leukemias

**DOI:** 10.1101/2023.03.16.532874

**Authors:** Leila Mahdavi, Haley Goodrow, Fatemeh Alikarami, Alexandra Lenard, Simone S. Riedel, Clara Libbrecht, Isabel Bowser, Sarah K. Tasian, Catherine D. Falkenstein, Bryan Manning, Sarah Skuli, Martin P. Carroll, Gerald Wertheim, Sheng F. Cai, Gerard McGeehan, Hongbo M. Xie, Kathrin M. Bernt

**Affiliations:** Division of Pediatric Oncology, Children’s Hospital of Philadelphia, Philadelphia, PA, USA; Department of Bioinformatics and Health Informatics (DBHI), Children’s Hospital of Philadelphia, Philadelphia, PA, USA; Department of Pediatrics, Perelman School of Medicine at the University of Pennsylvania and Abramson Cancer Center, Philadelphia, PA, USA; Department of Medicine, Perelman School of Medicine, University of Pennsylvania, PA 19104, USA; Department of Pathology, Children’s Hospital of Philadelphia, Philadelphia, PA, USA; Department of Medicine, Memorial Sloan Kettering Cancer Center, New York, NY, USA

## Abstract

Inhibitors of the Menin-KMT2A interaction are promising agents for the treatment of *KMT2A*-rearranged (*KMT2A*-r) leukemias. We evaluated Menin inhibition in patient derived xenografts of *KMT2A-r* leukemias with high-risk features. Three AMLs with high-risk fusion partners (*MLLT10, MLLT4*) and two infant ALL samples were sensitive to Menin inhibition. We also evaluated serial samples from two patients with multiply relapsed ALL. We found that highly pretreated *KMT2A-AFF1* ALL samples were much less sensitive compared to cells obtained earlier in the same patients’ disease course. Since none of the patients had been treated with a Menin inhibitor, resistance in these highly pretreated samples was acquired in the absence to Menin inhibitor exposure. Transcriptomic analysis documented sustained on-target efficacy towards the canonical targets in the Menin-inhibitor in resistant cells. Targeted genomic analysis documented the emergence of multiple co-mutations, including RAS pathway and *TP53* mutations, although neither was sufficient to induce Menin-inhibitor resistance in vitro. Downregulation of KMT3D may account for resistance in one patients; inactivation of KMT2C/D had previously been reported to result in Menin inhibitor resistance. Future studies will need to clarify more broadly which genomic/epigenomic alterations drive upfront resistance. Regardless of mechanism, our data supports using Menin-inhibitors upfront or in early lines of therapy before substantial genomic or epigenomic evolution has occurred.

## Introduction

10% of acute leukemias carry rearrangements of the *KMT2A* gene (Mixed-Lineage Leukemia-1, *MLL-1*). More than 80 fusion partners have been identified, and differences in fusion partner have been linked to outcomes (1-3). *KMT2A*-r leukemias have a unique gene expression profile characterized by high expression of HOXA cluster genes, Meis1 and FLT3 (4,5). The 5-year event free and overall survival of pediatric *KMT2A*-r AML remains suboptimal at 38% and 58% (3). The survival for infants and adults with *KMT2A*-r ALL remains dismal (6-8).

Recently, inhibition of the KMT2A-interacting protein Menin was identified as a promising therapeutic strategy (9-14). Menin is essential for the oncogenic activity of KMT2A fusions (15). Early studies of the Menin inhibitor VTP-50469 demonstrated impressive single agent activity in *KMT2A*-r ALL and AML (9,10,14). Molecularly, Menin inhibitors interfere with the recruitment of KMT2A-fusion to a subset of their target genes (9-11,13).

Preclinical studies published to date mostly interrogated common fusion partners and samples banked at initial diagnosis (9,10,12). We sought to evaluate the efficacy of Menin inhibition in AML with the more rare high-risk fusion partners *MLLT10 (AF10)* and *MLLT4 (AF6) (2,3)*, infant-ALL (6,7), and multiply relapsed *KMT2A*-r ALL (16). We found that highly pretreated ALL samples were less sensitive to Menin inhibition than samples from the same patients earlier in their course. Our data supports moving Menin-inhibitors into early lines of therapy to maximize their potential clinical impact.

## Materials and Methods

### Human samples

Samples were obtained from patients at the Children’s Hospital of Philadelphia with informed consent according to the Declaration of Helsinki and Institutional Review Board (IRB) approval.

### Patient derived xenografts

NSG (ALL samples, NOD-scid IL2Rgnull) or NSGS (AML samples, NOD-scid IL2Rgnull-3/GM/SF, Jackson laboratories®) were conditioned with Busulfan and patient cells were injected into the tail vein. Menin inhibitor or control chow was given as indicated. Human leukemia cells were detected in peripheral blood, bone marrow and spleen using anti-huCD45.

### RNA-Seq analysis

Raw Fastq files were aligned using STAR against reference Mus musculus GRCm38, read-counts were quantified by Kallisto and directly imported into DESeq2 (RRID:SCR_000154). Gene Set Enrichment Analysis (GSEA) was carried out using GSEA software (SeqGSEA, RRID:SCR_005724).

### Targeted DNA sequencing

Targeted sequencing of patient leukemia samples was performed using the in-house clinical platform. MEN1 mutation status was validated using clinical sequencing at Memorial Sloan Kettering Cancer Institute.

### Data availability statement

RNA-Seq data has been submitted to GEO and access will be provided upon request.

## Results

### Menin inhibition is effective in AML samples with high risk KMT2A fusion partners

We first evaluated the efficacy of Menin inhibition in two AML patient samples with the high-risk fusion partner MLLT-10 (AF10) (**Table 1** and **supplemental material)**. These experiments were carried out using the preclinical tool compound VTP50469. Menin inhibitor treated mice lived significantly longer than untreated mice. (**Figure 1A and supplementary Figure 1A**). Menin inhibition was also effective in a xenografts of a patient with high risk KMT2A-MLLT4 (AF6) AML (**supplementary Figure 2**), and 2 infant ALL samples at initial diagnosis (patient 4, 21). (**Figure 1B, and supplementary Figure 3**).

**Table 1:** Patient and sample characteristics. Detailed specimen reports, cytogenetic and molecular analysis and detailed clinical characteristics are available in the supplementary material. AML: acute myeloid leukemia, ALL: acute lymphoblastic leukemia, AUL: acute undifferentiated leukemia.

| # | Diagnosis | Age at diagnosis | Sex | Day from diagnosis | Prior lines of therapy | Fusion |
| --- | --- | --- | --- | --- | --- | --- |
| 1-1 | AML | 18 months | M | 0 | 0 | KMT2A-MLLT10 (MLL-AF10) |
| 2-1 | AML | 6.8 years | F | 0 | 0 | KMT2A-MLLT10 (MLL-AF10) |
| 3-1 | AUL | 12 years | M | 0 | 0 | KMT2A-MLLT4 (MLL-AF6) |
| 4-1 | Infant ALL | 10 months | F | 626 | 2 | KMT2A-MLLT6 (MLL-AF17) |
| 21-1 | Infant ALL | 1 months | F | 0 | 0 | KMT2A-AFF1 (MLL-AF4) |
| 5-1 | B-ALL | 13 months | M | 0 | 0 | KMT2A-AFF1 (MLL-AF4) |
| 5-2 | Lineage switch |  |  | 721 | 6 | KMT2A-AFF1 (MLL-AF4) |
| 8-1 | B-ALL | 14 years | F | 217 | 3 | KMT2A-AFF1 (MLL-AF4) |
| 8-2 | Lineage switch |  |  | 282 | 6 | KMT2A-AFF1 (MLL-AF4) |

**Figure 1:**
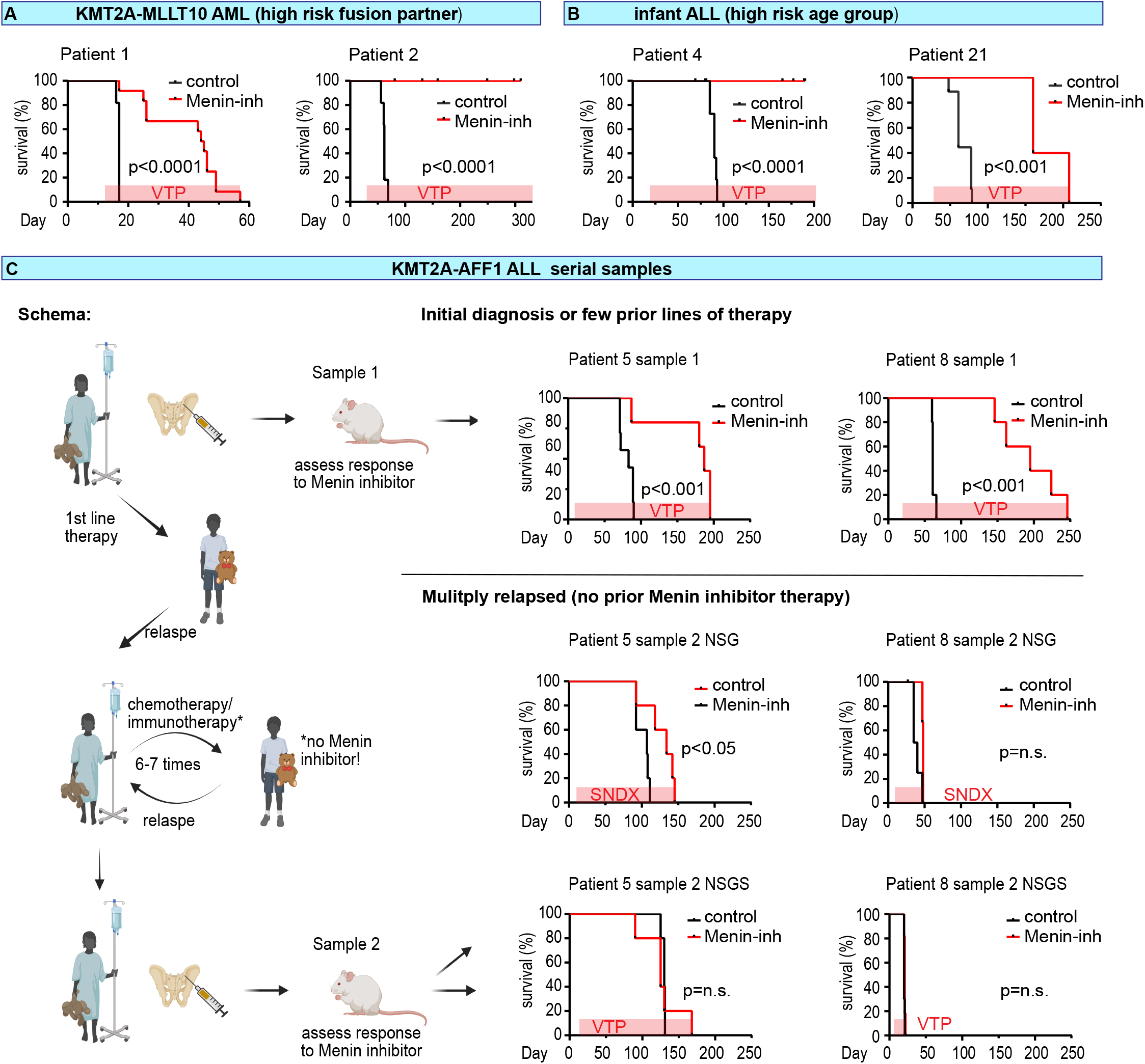
Menin-inhibitor treatment of high risk *KMT2A*-r leukemia xenografts. In vivo treatment of patient derived xenografts from *KMT2A* rearranged leukemias with Menin inhibitor (red) or control (black). The type of inhibitor indicated on the graph and choice of agent was based on availability. We found VTP-50469 and SNDX-5613 to be equipotent. **A:** AML patients with high-risk fusions partner *MLLT10, MLLT4*. **B**: infant B-ALL **C:** *KMT2A-AFF1* PDX established early (**top row**, initial diagnosis or three prior lines of therapy) or late (**middle and bottom row**, 6 prior lines of therapy) in the course of a patient’s therapy. Late samples were transplanted into either NSG (middle row) or NSGS (bottom row) mice. n=5-10 per treatment arm, error bars = SEM, *p<0.0001, Mantle-Cox.

### Preceding standard chemo / immunotherapy induces Menin-inhibitor resistance in multiply relapsed, Menin-inhibitor naïve *KMT2A*-r ALL samples

We next interrogated the efficacy of Menin inhibition in two paired samples from patients with *KMT2A-*r ALL, obtained early and late in their disease course. Due to availability, some of these experiments were carried out using the SNDX5613 compound. SNDX5613 is equipotent to VTP50469 in our xenograft model (**Supplementary Figure 4**). “Early” samples were from either initial diagnosis (patient 5) or the third relapse (patient 8). “Late” samples were obtained after several lines of therapy, including high dose chemotherapy, immunotherapy and stem cell transplant. Patient 5 and 8 experienced a lineage switch to AML (for detail see supplemental material). Menin inhibition has been shown to induce myeloid differentiation in AML samples, which could be promoted by the presence of myeloid cytokines. In order to facilitate a differentiation response in lineage switch samples, “late” samples were transplanted into NSGS mice (which express human cytokines that promote differentiation, including interleukin 3 and GM-CSF), as well as NSG mice, while initial ALL samples were only transplanted into NSG mice. We found that leukemia cells obtained early in the disease course were sensitive to Menin inhibition (**Figure 1C top row**). In contrast, leukemia samples obtained after multiple lines of therapy from patients 5 and 21 were near completely resistant (**Figure 1C middle and bottom row, and supplementary Figure 5 - 7**). Menin inhibitor resistance thus had developed without exposure to Menin inhibition.

### RAS pathway and *TP53* mutations do not mediate Menin-inhibitor resistance

We next analyzed the co-mutational landscape of our serial samples using a targeted DNA sequencing panel (**Figure 2A**). We found RAS pathway mutations in all three, and *TP53* mutations in two of three patients. We did not observe any of the mutations in the *MEN1* gene (targeted sequencing and RNA-Seq) (17). Prior reports have documented that *KMT2A*-fusion leukemias carrying *TP53* mutations or activating RAS mutations can be highly sensitive to Menin inhibition (18-20). We confirmed these observations by treating a murine model of *KMT2A-MLLT2* AML with and without NRAS mutation, (**Figure 2B**), and MV;11 cells engineered to carry inactivating mutations of *TP53* to Menin-inhibition (**Figure 2C**). Neither *NRAS* nor *TP53* mutations induced Menin inhibitor resistance.

**Figure 2:**
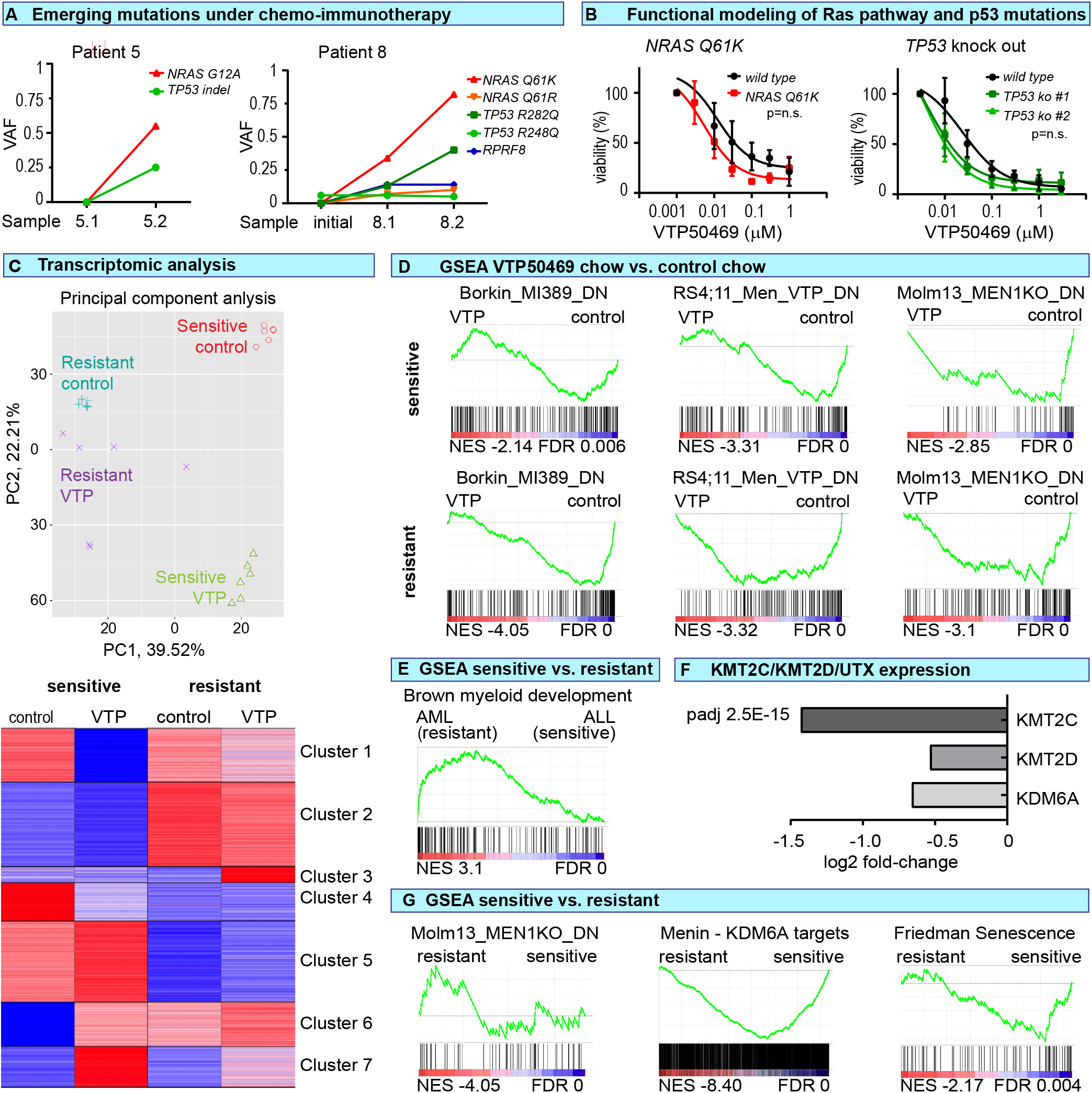
molecular characterization of patient samples over the course of preceding therapy. **A:** Variant allelic frequency (VAF) of mutations identified on targeted NGS of primary patient bone marrow cells obtained at different time points during the treatment course corresponding to, or close to, the time points from which PDXs were established (left: patient 5, right: patient 8). **B:** Functional modeling of RAS pathway and *TP53* mutations. **Left:** in vitro treatment response of murine KMT2A-MLLT3 driven AML cells with or without expression of NRAS-Q61K (retroviral model) to VTP50469. **Right:** in vitro treatment response of MV4;11 (*KMT2A-AFF1, TP53* wild type) as well as CRISPR engineered biallelic *TP53* mutant sublines (*TP53*-KO1 and 2) to VTP50469. Cell viability after 96 hours of drug treatment was determined by trypan blue exclusion. N=3, error bar = SEM, differences between WT and KO were not statistically significant. **C-F:** RNA-Seq of sorted leukemia cells (huCD45+, mCD45-, DAPI-) from the bone marrow of mice treated with VTP-50469 for 3 weeks (Sample 8-1 = sensitive, sample 8-2 = resistant). **C:** Principal component analysis and K-means clustering. **D:** GSEA of established Menin-response signatures. Top row: sensitive, second row: resistant samples. Gene signatures contain genes that are downregulated upon Menin inhibitor treatment in MV4;11 cells (left panels), sites that lose Menin binding and are downregulated upon Menin inhibitor treatment in RS4;11 cells (middle panels), and genes that are downregulated upon Menin deletion in Molm13 cells (right panels). **E-G:** Comparison of resistant and sensitive samples isolated from animals in the vehicle control group (i.e. no Menin inhibitor exposure). **E**: enrichment of myeloid developmental genes in the lineage switch resistant sample. **F**: downregulation of KMT2C in the resistant sample (p=2.5E-15). KMT2D and the core complex member KDM6A (UTX) are also significantly downregulated (*padj<0.05). **G:** Comparison of resistant and sensitive samples isolated from animals in the Menin inhibitor treatment group. Left panel: no enrichment of core Menin-KMT2A-fusion program. Middle panel: enrichment of KDM6A dependent genes in sensitive cells (Menin-UTX Targets). Right panel: enrichment of senescence associated genes in sensitive cells.

### Sustained suppression of the canonical Menin:KMT2A-fusion targets in resistant cells

We next performed RNA-Seq of leukemia isolated from xenografts, comparing the transcriptional response to *in vivo* Menin inhibition in samples from patient 8 obtained early (sample 8-1, “sensitive”) and late (sample 8-2, “resistant”) in the disease course. Mice were treated for 3 weeks with Menin inhibitor or control, then leukemia cells were isolated and subjected to RNA-Seq. Principal component analysis demonstrated that Menin inhibition induced a stronger transcriptional response in the samples obtained earlier in the disease course (**Figure 4A**). However, and somewhat unexpectedly, profound downregulation of the canonical Menin-inhibitor responsive gene signature was maintained in the resistant AML (**Figure 4B, Supplementary Figure 6, Supplementarry Table 1**). Enriched signatures included genes downregulated upon Menin inhibitor treatment (13), sites that lose Menin binding and are downregulated upon Menin inhibitor treatment (10), genes downregulated upon Menin deletion (10), as well as genes regulated by HOXA9 (25). We conclude that Menin inhibition maintained on-target transcriptional activity in resistant cells.

### Decreased expression of KMT2C and decreased KMT2C/KDM6A target induction in multiply relapsed cells

We next compared sensitive and resistant cells. Consistent with the clinically observed myeloid switch we observed enrichment of a myeloid differentiation signature in the resistant cells (26) (**Figure 4C**). A pattern of Menin-inhibitor resistance with preserved transcriptional response of the core Menin:KMT2A-fusion target genes that emerged under pressure from Menin-inhibitor exposure was recently reported. Resistance in this context was linked to a blunted induction of a non-canonical senescence signature that requires KMT2C/KDM6A (UTX) (24). We therefore probed the expression of KMT2C/D complex members and found profound downregulation of KMT2C in resistant cells (**Figure 4D)**. The core Menin:KMT2A-fusion target signature did not enrich in either sensitive or resistant cells (**Figure 4E**) (10). However, KDM6A-targets (24) and senescence associated genes (27) were enriched in sensitive cells from the VTP treatment group. The downregulation of KMT2C in resistant cells was therefore associated with a blunted induction of the KMT2D/KDM6A dependent senescence signature described by Soto-Feliciano and colleagues (24), and could potentially account for the decreased sensitivity to Menin inhibition.

## Discussion

This is the first report showing that preceding chemo/immunotherapy can render *KMT2A*-rearranged samples resistant to Menin inhibition without prior Menin inhibitor exposure. Our findings complement several recent reports of resistance that arises under pressure of Menin inhibitor treatment.

This study has potential clinical implications. The majority of patients in the recent phase I / II study of revumenib (SNDX-5613) saw downregulation of the key KMT2A target genes. The overall response and CR/CRh rates, however, were much lower at 50% and 30%(14). While this is an excellent CR rate for a single agent in a multiply relapsed, highly refractory patient population, our data suggest that an even higher percentage of patients might have responded if treated in an earlier line of therapy.

On a mechanistic level, we found that the transcriptional response of the core Menin-KMT2A-fusion target genes was preserved in resistant cells. However, resistant cells had become tolerant to the downregulation of key KMT2A fusion target genes over the course of preceding chemo/immunotherapy. While two patients (Pt 5 and 8) underwent a lineage switch (28), myeloid differentiation alone seems unlikely to mediate resistance given the encouraging clinical response rates in AML (29). However, failure to induce a KMT2C/KDM6A dependent senescence signature, a described mechanism of Menin inhibitor resistance (24), was observed in one patient. Our data suggests that KMT2C/KDM6A might be involved in mediating resistance both upfront and in response to Menin inhibitor treatment.

## Conclusion

We document the emergence of Menin inhibitor resistance in highly pretreated, Menin-inhibitor naïve, ALL samples that were initially sensitive. (17). Our data supports treating patient with Menin-inhibitors early in their disease course before significant resistance has developed.

## Acknowledgments

We thank patients and families for providing samples for this study. We thank Florin Tuluc and the staff at the Children’s Hospital of Philadelphia flow core for assistance with cell sorting. We thank all of the veterinary staff at the Children’s Hospital of Philadelphia for assistance in caring for the mice. This work was supported by funding from the Children’s Hospital of Philadelphia Research Institute [to KMB], the Emerson Collective [to KMB], and gifts from “Association Raphael” and “Association 111 des arts, Lyon” [to CL] and the SchylerStrong Foundation [to SKT]. SKT is a Scholar of the Leukemia & Lymphoma Society and holds the Joshua Kahan Endowed Chair in Pediatric Leukemia Research at the Children’s Hospital of Philadelphia. Syndax provided VTP/SNDX-chow as well as limited funding for in vivo experiments [to KMB].

