## Supplemental Figures, Materials and Methods for "Upfront Menin-inhibitor resistance in multiply pretreated leukemias"

**SUPPLEMENTARY DATA**

1. **Supplementary figures 1-6**
2. **Patient specimen reports**
3. **Patient vignettes**
4. **Supplementary materials and methods**
5. **Supplementary references**

**Two supplementary tables are provided as separate files.**

**A:** PDX1 (*KMT2A-MLLT10* AML)

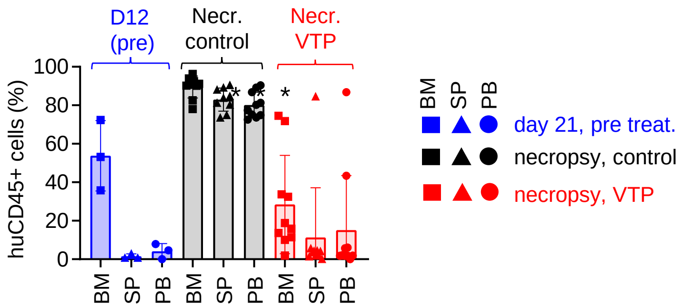


**B**: PDX2 (*KMT2A-MLLT10 AML*)

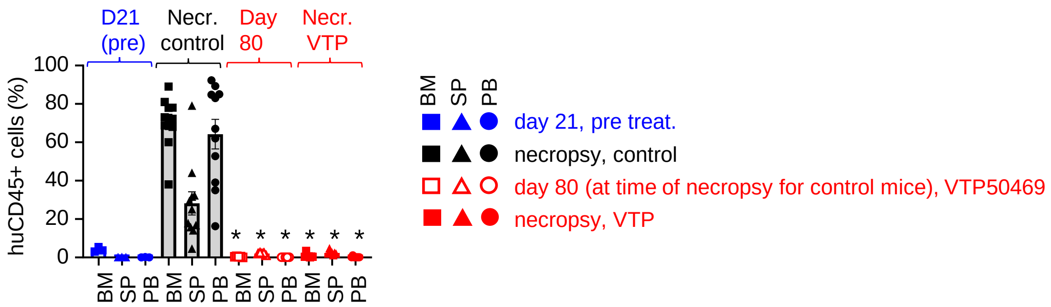


**Supplementary Figure 1:** In vivo treatment of patient derived xenografts from *KMT2A-MLLT10* rearranged acute myeloid leukemias with the Menin inhibitor (VTP-50469, red) or control (black). Blue denotes pre-treatment time points.
Leukemic burden in bone marrow (BM), spleen (SP) and peripheral blood (PB) (measured as huCD45+/mCD45-) at the indicated time points. **A:** PDX1. Disease evaluation at necropsy shows animals succumbed to leukemia. This model causes CNS disease, some of the treated mice with low leukemic burden at necropsy presented with hind limb paralysis.
**B:** PDX 2. 5 surviving VTP treated mice were sacrificed on day 80, when all control mice had succumbed to disease. Evaluation revealed no evidence of leukemia (day 80, open symbols). The remaining 6 animals were observed for survival analysis and eventually reached the end of their natural life span without evidence of disease.

n=3 pre treatment, n=5 Pt 2 day 80, n=5-10 at necropsy per treatment arm and time point error bars = SEM, *p<0.0001, unpaired t-test (comparison between Menin inhibitor and control within the same tissue).

PDX2 (*KMT2A-MLLT4* AML)

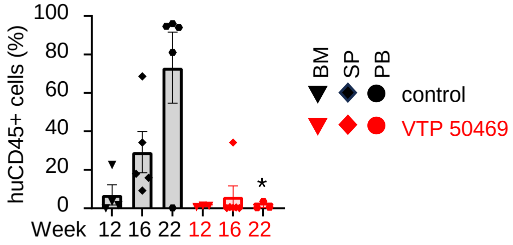


**Supplementary Figure 2:** In vivo treatment of patient derived xenografts from *KMT2A-MLLT4 (MLL-AF6)* rearranged AML with the Menin inhibitor (VTP-50469, red) or control (black). Shown is leukemic burden (right) in bone marrow (measured as huCD45+/mCD45-) at the indicated time points. Due to the pandemic lock down this experiment was terminated at week 22 and only limited analysis was obtained (bone marrow flowcytometry only, no spleen or peripheral blood analysis). Survival was not obtained as an endpoint.

n= 5 per treatment arm and time point, statistical analysis: unpaired t-test at each time point, error bars = SEM, $^*p<0.001$.

**A:** PDX4 (*KMT2A-MLLT6* infant ALL)

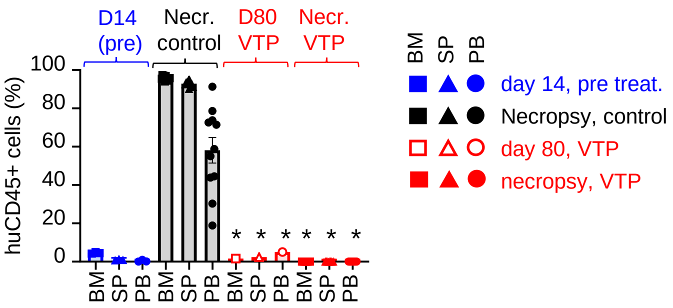


**B**: PDX21 (*KMT2A-AFF1* infant ALL)

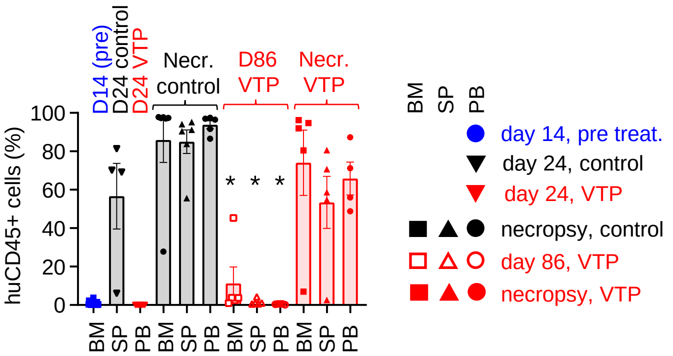


**Supplementary Figure 3:** In vivo treatment of patient derived xenografts from *KMT2A* rearranged infant leukemias (B-ALL) with the Menin inhibitor (VTP-50469, red) or control (black). Blue denotes pre-treatment time points. Leukemic burden in bone marrow (BM), spleen (SP) and peripheral blood (PB) (measured as huCD45+/mCD45-) at the indicated time points.

**A:** PDX4 (*KMT2A-MLLT6)* infant ALL. 5 surviving VTP treated mice were sacrificed on day 80, when all control mice had succumbed to disease. Evaluation revealed no evidence of leukemia (day 80, open symbols). The remaining 5 animals were observed for survival analysis and eventually reached the end of their natural life span without evidence of disease.

**B:** PDX 21 *(KMT2A-AFF1*) infant ALL. Due to limited animal numbers, leukemic burden was assessed in the blood on day 14 (start of treatment), but not animals were sacrificed at the pre-treatment time point. Blood leukemic burden was assessed again on day 24. On day 86, when all control mice had succumbed to disease, 5 Menin-inhibitor treated mice were sacrificed. Evaluation revealed very low or no evidence of leukemia (day 86, open symbols). The remaining 5 animals were observed for survival analysis and eventually succumbed to disease

n=9-10 per treatment arm, error bars = SEM, *p<0.0001, unpaired t-test (comparison between Menin inhibitor and control within the same tissue).

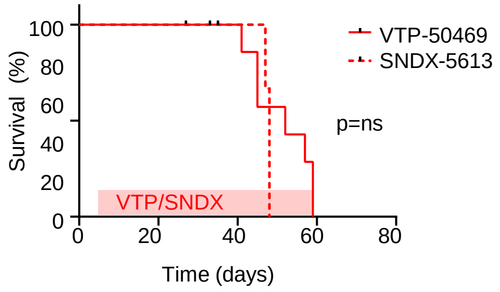


**Supplementary Figure 4: Similar efficacy of VTP-50469 and SNDX-5613 in NSG mice.**
In vivo treatment of a *KMT2A* rearranged leukemia patient derived xenografts the Menin inhibitor VTP-50469 (solid line) or the equipotent SNDX-5613 (dashed line). 10,000 cells per mouse, n=5-7 per treatment arm, *p=not significant (ns), Mantle-Cox

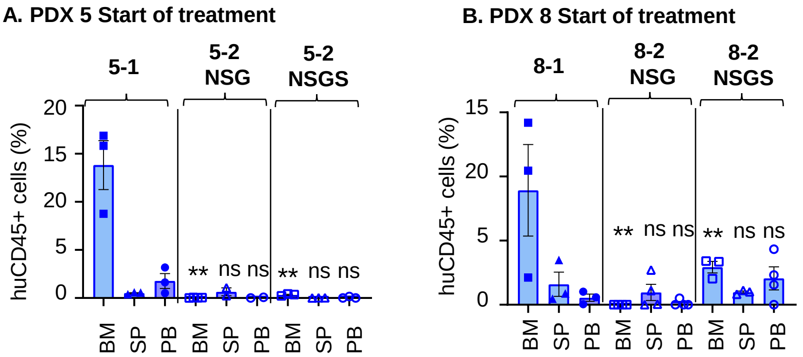


**Supplementary Figure 5: Leukemic burden at the start of treatment for serial ALL samples.**

Leukemic burden at the start of treatment in bone marrow (BM), spleen (SP) and peripheral blood (PB) measured as huCD45+/mCD45-. n=3, error bars = SEM, **p<0.01, unpaired t-test (comparison between Menin inhibitor and control within the same tissue).

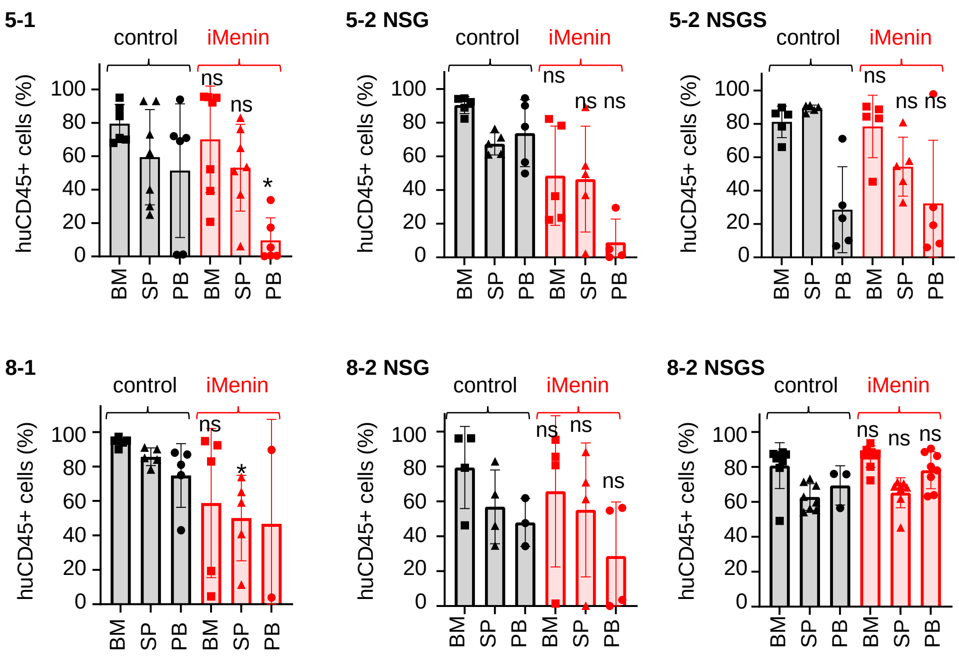


**Supplementary Figure 6: Leukemic burden at necropsy**

Leukemic burden measured as huCD45+/mCD45- in in bone marrow (BM), spleen (SP) and peripheral blood (PB) at necropsy.

n=5-10 per treatment arm, error bars = SEM, *p<0.05,**p<0.01, unpaired t-test (comparison between Menin inhibitor and control within the same tissue).

**Survival in NSG mice - secondary transplant from (A):**

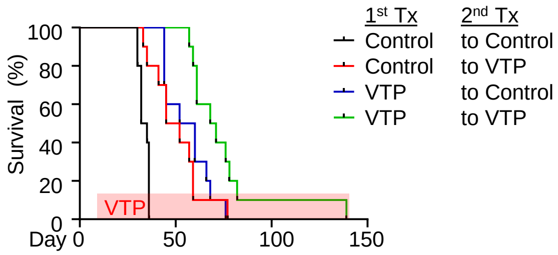


**Supplementary Figure 7: Re-transplantation of Menin inhibitor treated cells (PDX 8-2).**

**Secondary** transplantation of 10,000 cells from the primary transplant in Figure 1 (Patient 8 sample 2 NSGS) into NSG mice. Cells from both VTP-50469 and control arms were each transplanted into secondary cohorts that were assigned to VTP and control respectively.
Start of inhibitor treatment day 7. Overall survival, n=10 per treatment arm.
***p<0.001, Mantle-Cox (survival, pairwise).
All treatment arms were statistically significant compared to control arm (control-control) at p<0.001, documenting that VTP-50469 still retains some activity at this time point. VTP-VTP was statistically different from all other arms at p<0.001.
Despite transplantation of very low cell numbers and extending the treatment period through re-transplantation, the difference between treatment and control arms is still much smaller than in sample 8-1 earlier in the patient's disease course.

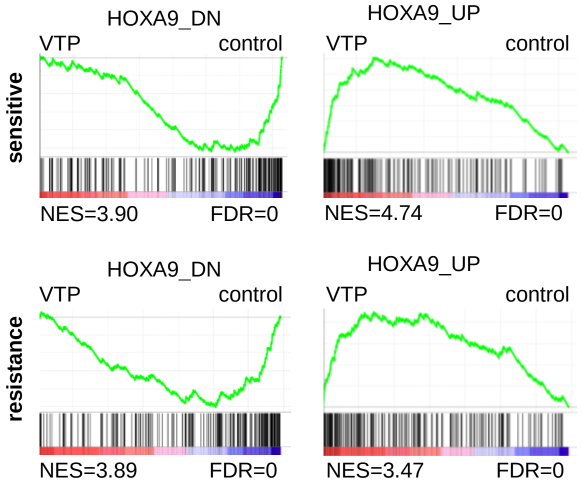


**Supplementary Figure 6**

GSEA of HOXA9 dependent gene signatures in VTP versus control treated cells. Top row: sensitive, second row: resistant samples.

Gene signatures contain genes that are downregulated upon knockdown of HOXA9 (left panels) and upregulated upon knockdown of HOXA9 (right panels)

### SPECIMEN REPORTS

**Sample ID: PDX-1-1**
Gender: Male
Race: White
Cancer Type: AML
CNS Status: CNS1
Age at diagnosis: 18 months
Cause of Death: r/r AML

**Cytogenetics**: 46,XY,?t(10;11)(p12;q23)[2]/46,XY[17

#### Fusions:

| *fusion* | *5_fusion transcript* | *3_fusion transcript* |
|---|---|---|
| *KMT2A-MLLT10* | KMT2A (NM_005933.3) exon 8 | MLLT10 (NM_004641.3) exon 10 |

#### Variants:

| *gene_name* | *position* | *assembly* | *isoform* | *nucleotide* | *amino_acid* | *vaf* |
|---|---|---|---|---|---|---|
| *CREBBP* | chr16:3778439- 3778456 | HG19 | NM_004380.2 | c.6600 6608delGCAGCAGCA | p.Gln2214 Gln2216del | 0.5 |
| *CREBBP* | chr16:3778448- 3778449 | HG19 | NM_004380.2 | c.6599 6600insACA | p.Gln2216dup | 0.21 |
| *CSF1R* | chr5:149441342- 149441342 | HG19 | NM_005211.3 | c.1697C>G | p.Pro566Arg | 0.05 |
| *NRAS* | chr1:115258748- 115258A748 | HG19 | NM_002524.4 | c.34G>& | p.Gly12Cys | 0.17 |
| *NSD2* | chr4:1936899- 1936899 | HG19 | NM_133335.3 | c.1584C>A | p.His528Gln | 1.43 |
| *TERT* | chr5:1254594- 1254594 | HG19 | NM_198253.2 | c.3184G>A | p.Ala1062Thr | 0.49 |

**Sample ID: PDX-2-1**
Gender: Female
Race: White
Cancer Type: AML
Age at diagnosis: 6.8 years
CNS Status: CNS1

**Cytogenetics**:
47,XX,+4,der(10)t(10;11)(p12;q23)inv(11)(q21q23),der(11)t(10;11)(p12;q21),der(22

### Fusions:

| *fusion* | *5_fusion transcript* | *3_fusion transcript* |
|---|---|---|
| *KMT2A-MLLT10* | KMT2A (NM_005933.3) exon 9 | MLLT10 (NM_004641.3) exon 9 |

### Variants:

| *gene_name* | *position* | *assembly* | *isoform* | *nucleotide* | *amino_acid* | *vaf* |
|---|---|---|---|---|---|---|
| *KMT2D* | chr12:49448143-49448143 | HG19 | NM_003482.3 | c.457G>A | p.Glu153Lys | 0.5 |
| *NRAS* | chr1:115256528-115256528 | HG19 | NM_002524.4 | c.183A>C | p.Gln61His | 0.41 |
| *SETD2* | chr3:47108609-47108609 | HG19 | NM_014159.6 | c.6061-1G>A | | 0.13 |
| *SETD2* | chr3:47155457-47155457 | HG19 | NM_014159.6 | c.4624A>G | p.Arg1542Gly | 0.44 |

**Sample ID: PDX-3-1**
Gender: Male
Race: White
Cancer Type: AUL
Age at diagnosis: 12 years
CNS Status: CNS1

**Cytogenetics**: 46,XY,der(6)(6pter->6q2?6::?),der(11)(11pter->11q23::6q27->6qter),del(11)(q2?2q2 ?3),der(17)(11qter->11q23::17p13->17q2?5::11q2?2->q2?3::17q2?5->17qter)[20].ish der(11(11ptel+,5'KMT2A+,3'KMT2A-,6qtel+,11qtel-),del(11)(11ptel+,KMT2A-,11qtel+ ), der(17)(17ptel+,CEP17+,17qtel+, 5'KMT2Ax1,3'KMT2Ax2)(5'KMT2A con 3'KMT2A2x1)

**Fusions:**

| *fusion* | *5_fusion transcript* | *3_fusion transcript* |
|---|---|---|
| *KMT2A-MLLT4* | KMT2A (NM_005933.3) exon 10 | MLLT4 NM_001040000.2) exon 2 |

**Variants:**

| *gene_name* | *position* | *assembly* | *isoform* | *nucleotide* | *amino_acid* | *vaf* |
|---|---|---|---|---|---|---|
| *ASXL1* | chr20:31022441-31022442 | HG19 | NM_015338.5 | c.1934dupG | p.Gly646Trpfs*12 | 0.4 |
| *BCORL1* | chrX:129155008-129155009 | HG19 | NM_021946.4 | c.3496dup | p.Ala1166Glyfs*50 | 0.06 |
| *FLT3* | chr13:28608257-28608258 | HG19 | NM_004119.2 | c.1798_1799 insGGATCCCCGATTTCAGAGAATATGAATATG | p.Tyr599_Asp600 insGlyIleProAspPheArgGluTyrGluTyr | 0.15 |
| *PHF6* | chrX:133549142-133549143 | HG19 | NM_032458.2 | c.826delinsGGGCGT | p.Lys276Glyfs*5 | 0.99 |
| *RUNX1* | chr21:36171615-36171625 | HG19 | NM_001754.4 | c.940_950delinsCCA | p.Ser314Profs*283 | 0.3 |
| *SUZ12* | chr17:30322767-30322768 | HG19 | NM_015355.3 | c.1786dup | p.Thr596Asnfs*6 | 0.08 |
| *USH2A* | chr1:216462627-216462627 | | NM_206933.2 | c.1966G>A | p.Asp656Asn | 0.51 |

**Sample ID: PDX-4-1**
Gender: Female
Race: White
Cancer Type: relapsed Infant ALL (late relapse)
Age at diagnosis: 10 months
CNS Status: CNS2a

**Cytogenetics (initial diagnosis)**: 46,XX[19]

**Cytogenetics (sample)**: 49,XX,+X,+6,t(11;17)(q23;q12),+21[11]//46,XY[9]

**Fusions:**

| *fusion* | *5_fusion transcript* | *3_fusion transcript* |
|---|---|---|
| *KMT2A-MLLT6* | KMT2A (NM_005933.3) exon 10 | MLLT6 (NM_005937.3) exon 7 |
| *KMT2A-MLLT6* | KMT2A (NM_005933.3) exon 9 | MLLT6 (NM_005937.3) exon 7 |

**Variants (initial diagnosis):**

| *gene_name* | *position* | *assembly* | *isoform* | *nucleotide* | *amino_acid* | *vaf* |
|---|---|---|---|---|---|---|
| *SETBP1* | chr18:42643559-42643559 | HG19 | NM_015559.2 | c.4687C>A | p.Pro1563Thr | 0.47 |

**Variants (sample):**

| *gene_name* | *position* | *assembly* | *isoform* | *nucleotide* | *amino_acid* | *vaf* |
|---|---|---|---|---|---|---|
| *SETBP1* | chr18:42643559-42643559 | HG19 | NM_015559.2 | c.4687C>A | p.Pro1563Thr | 0.55 |

**Sample ID: PDX-5-1**
Gender: Male
Race: White
Cancer Type: B-ALL (initial diagnosis)
Cancer Risk: HR
Age at diagnosis: 13 months
CNS Status: CNS3
Cause of Death: r/r AML

**Cytogenetics (initial diagnosis/sample)**: 40~46,XY,t(4;11)(q21;q23)[cp14]/46,XY[1]

**Fusions:**

| *fusion* | *5_fusion transcript* | *3_fusion transcript* |
|---|---|---|
| *KMT2A-AFF1* | KMT2A (NM_005933.3) exon 8 | AFF1 (NM_005935.3) exon 4 |

**Variants (initial diagnosis/sample):**

| *gene_name* | *position* | *assembly* | *isoform* | *nucleotide* | *amino_acid* | *vaf* |
|---|---|---|---|---|---|---|
| *BCL6* | chr3:187446313-187446313 | HG19 | NM_001706.4 | c.1275C>T | p.Arg459Cys | 0.41 |

Copy number variant (initial diagnosis): loss of chromosome 7p (*including IKZF1*)

**Sample ID: PDX-5-2**
Gender: Male
Race: White
Cancer Type: B-ALL (relapsed)
Cancer Risk: HR
Age at diagnosis: 13 months
CNS Status: CNS3
Cause of Death: r/r lineage switch AML

**Cytogenetics (initial diagnosis)**: 40~46,XY,t(4;11)(q21;q23)[cp14]/46,XY[1]
**Cytogenetics (first relapse)**: 47,XY,t(4;11)(q21;q23),+8[17]

**Fusions (present at all time points):**

| *fusion* | *5_fusion transcript* | *3_fusion transcript* |
|---|---|---|
| *KMT2A-AFF1* | KMT2A (NM_005933.3) exon 8 | AFF1 (NM_005935.3) exon 4 |

**Variants (initial diagnosis):**

| *gene_name* | *position* | *assembly* | *isoform* | *nucleotide* | *amino_acid* | *vaf* |
|---|---|---|---|---|---|---|
| *BCL6* | chr3:187446313-187446313 | HG19 | NM_001706.4 | c.1275C>T | p.Arg459Cys | 0.41 |

**Variants (first relapse):**

| *gene_name* | *position* | *assembly* | *isoform* | *nucleotide* | *amino_acid* | *vaf* |
|---|---|---|---|---|---|---|
| *NRAS* | chr1:115258747-115258747 | HG19 | NM_002524.4 | c.35G>C | p.Gly12Ala | 0.32 |
| *TP53* | chr17:7577094-7577095 | HG19 | NM_000546.5 | c.844delinsTCCGG | p.Arg282Serfs*25 | 73 |
| *BCL6* | chr3:187446313-187446313 | HG19 | NM_001706.4 | c.1275C>T | p.Arg459Cys | 0.44 |

**Variants (last time point prior to sample time point):**

| *gene_name* | *position* | *assembly* | *isoform* | *nucleotide* | *amino_acid* | *vaf* |
|---|---|---|---|---|---|---|
| *NRAS* | chr1:115258747-115258747 | HG19 | NM_002524.4 | c.35G>C | p.Gly12Ala | 0.05* |
| *TP53* | chr17:7577094-7577095 | HG19 | NM_000546.5 | c.844delinsTCCGG | p.Arg282Serfs*25 | 0.11* |
| *BCL6* | chr3:187446313-187446313 | HG19 | NM_001706.4 | c.1275C>T | p.Arg459Cys | 0.51 |

*adjusted for bone marrow infiltration at 20%:
calculated VAF for NRAS: 0.25%,
calculated VAF for TP53: 55%

**Copy number variant (initial diagnosis):** loss of chromosome 7p
**Copy number variant (first relapse):** loss of chromosome 7p, gain of chromosome 8, loss of chromosome 17p (including PRPF8, TP53)
**Copy number variant (last time point prior to sample time point):** loss of chromosome 17p (including PRPF8, TP53)

**Sample ID: PDX-8-1**
Gender: Female
Race: White
Cancer Type: B-ALL
Cancer Risk: HR
CNS Status: CNS2a
Cause of Death: r/r lineage switch AML

**Specimen pathology report:** B-ALL

**FISH (initial diagnosis):** break apart probe set detected a split of 5'KMT2A signal from the 3'KMT2A signal in 97.5% (195/200) of the nuclei examined. Cytogenetics failed

**Cytogenetics (specimen):**
47,XX,t(4;11)(q21;q23),+der(4)t(4;11),der(17)t(8;17)(q13;p13)[12]/50~51,idem,+der(4)t(4;11),+21,+21[cp3]/46,XX[4]

**Fusions (initial diagnosis and specimen):**

| *fusion* | *5_fusion transcript* | *3_fusion transcript* |
|---|---|---|
| *KMT2A-AFF1* | KMT2A (NM_005933.3) exon 8 | AFF1 (NM_005935.3) exon 4 |

**Variants (initial diagnosis):**

| *gene_name* | *position* | *assembly* | *isoform* | *nucleotide* | *amino_acid* | *vaf* |
|---|---|---|---|---|---|---|
| *BCORL1* | chrX:129149813-129149813 | HG19 | NM_021946.4 | c.3065T>C | p.Ile1022Thr | 0.52 |
| *NOTCH1* | chr9:139409976-139409976 | HG19 | NM_017617.4 | c.1862G>A | p.Arg621His | 0.51 |
| *PAX5* | chr9:36882049-36882049 | HG19 | NM_016734.2 | c.964G>A | p.Ala322Thr | 0.5 |

**Variants (specimen):**

| *gene_name* | *position* | *assembly* | *isoform* | *nucleotide* | *amino_acid* | *vaf* |
|---|---|---|---|---|---|---|
| *BCORL1* | chrX:129149813-129149813 | HG19 | NM_021946.4 | c.3065T>C | p.Ile1022Thr | 0.52 |
| *NOTCH1* | chr9:139409976-139409976 | HG19 | NM_017617.4 | c.1862G>A | p.Arg621His | 0.51 |
| *NRAS* | chr1:115256529-115256529 | HG19 | NM_002524.4 | c.182A>G | p.Gln61Arg | 0.06 |
| *NRAS* | chr1:115256530-115256530 | HG19 | NM_002524.4 | c.181C>A | p.Gln61Lys | 0.13 |
| *PRPF8* | chr17:1557248-1557248 | HG19 | NM_006445.3 | .6050C>T | p.Ser2017Leu | 0.14 |
| *TP53* | chr17:7577093-7577093 | HG19 | NM_000546.5 | c.845G>C | p.Arg282Pro | 0.34 |
| *TP53* | chr17:7577538-7577538 | HG19 | NM_000546.5 | c.743G>A | p.Arg248Gln | 0.07 |

**Sample ID: PDX-8-2**
Gender: Female
Race: White
Cancer Type: B-ALL with lineage switch
Cancer Risk: HR
CNS Status: CNS2a
Cause of Death: r/r lineage switch AML

**Specimen pathology report:** AML (B-ALL w Lineage Switch)

**Cytogenetics - specimen:**
48,XX,t(4;11)(q21;q23),+der(4)t(4;11),+6,der(17)t(8;17)(q13;p13)[17]/52,idem,+5,+9,+21,+21 [1]

**Fusion:**

| *fusion* | *5_fusion transcript* | *3_fusion transcript* |
|---|---|---|
| *KMT2A-AFF1* | KMT2A (NM_005933.3) exon 8 | AFF1 (NM_005935.3) exon 4 |

**Variants:**

| *gene_name* | *position* | *assembly* | *isoform* | *nucleotide* | *amino_acid* | *vaf* |
|---|---|---|---|---|---|---|
| *BCORL1* | chrX:129149813-129149813 | HG19 | NM_021946.4 | c.3065T>C | p.Ile1022Thr | 0.52 |
| *NOTCH1* | chr9:139409976-139409976 | HG19 | NM_017617.4 | c.1862G>A | p.Arg621His | 0.51 |
| *NRAS* | chr1:115256529-115256529 | HG19 | NM_002524.4 | c.182A>G | p.Gln61Arg | 0.05 |
| *NRAS* | chr1:115256530-115256530 | HG19 | NM_002524.4 | c.181C>A | p.Gln61Lys | 0.4 |
| *PRPF8* | chr17:1557248-1557248 | HG19 | NM_006445.3 | .6050C>T | p.Ser2017Leu | 0.14 |
| *TP53* | chr17:7577093-7577093 | HG19 | NM_000546.5 | c.845G>C | p.Arg282Pro | 0.82 |
| *TP53* | chr17:7577538-7577538 | HG19 | NM_000546.5 | c.743G>A | p.Arg248Gln | 0.10 |

**Sample ID: PDX-21-1**
Gender: Female
Race: White
Cancer Type: B-ALL
Cancer Risk: HR
CNS Status: CNS2c
Cause of Death: r/r ALL

**Specimen pathology report:** B-ALL

**Cytogenetics:** 46,XX,t(4;11)(q21;q23)[20]

**Fusion:**

| *fusion* | *5_fusion transcript* | *3_fusion transcript* |
|---|---|---|
| *KMT2A-AFF1* | KMT2A (NM_005933.3) exon 10 | AFF1 (NM_005935.3) exon 4 |

**Variants:**

| *gene_name* | *position* | *assembly* | *isoform* | *nucleotide* | *amino_acid* | *VAF* |
|---|---|---|---|---|---|---|
| *DDX41* | chr5:176942773 | HG19 | NM_016222.3 | c.484C>T | p.Arg162Cys | 0.47 |
| *KMT2D* | chr12:49421617 | HG19 | NM_003482.3 | c.14612G>A | p.Ser4871Asn | 0.22 |
| *KRAS* | chr12:25398281 | HG19 | NM_033360.3 | | p.Gly13Asp | 0.14 |
| *NRAS* | chr1:115258744 | HG19 | NM_002524.4 | c.38G>A | p.Gly13Asp | 0.13 |
| *NT5C2* | chr10:104851320-104851321 | HG19 | NM_012229.4 | c.1211del | p.Lys404Serfs*23 | 0.45 |

**Patient Vignette for PDX 8-1 (ALL2184) and 8-2 (AML2263) primary samples and PDX models[1],[2]**

A 13 year-old female was diagnosed with CD19+ NCI HR B-acute lymphoblastic leukemia (B-ALL) with WBC of > 450,000 with 95% peripheral blasts. Cerebrospinal fluid (CSF) had microscopic evidence of leukemia (CNS2a). Initial cytogenetic analysis failed. Fluorescence in situ hybridization (FISH) assays detected KMT2A rearrangement in 97.5% of cells. RNA-based fusion and DNA-based next-generation sequencing (NGS) analyses demonstrated KMT2A::AFF1 rearrangement and no known oncogenic point mutations or indels (the observed variants of BCORL1, NOTCH1 and PAX5 were interpreted as likely constitutional and of unknown significance). The patient received four-drug induction chemotherapy as per the Children's Oncology Group (COG) AALL1131 study and cleared her CSF, but had positive end-induction measurable residual disease (MRD) at 0.77% in her bone marrow. She received consolidation chemotherapy as per AALL1131 very high-risk (VHR) arm A and had positive end-consolidation MRD at 0.19%. She underwent autologous T cell apheresis for future CD19 chimeric antigen receptor (CAR) T cell immunotherapy (tisagenlecleucel). She experienced a frank relapse (#1) that was treated with inotuzumab, but again had progressive B-ALL in peripheral blood 7 months after her initial diagnosis. Repeat genetic analysis redetected the known KMT2A::AFF1 fusion and acquisition of new somatic NRAS (Q61R with variant allele frequency [VAF] 0.06, Q61K with VAF 0.13) and TP53 (R248Q with VAF 0.0.07, R248P with VAF 0.34) mutations (specimen time point for PDX8-1, after 3 lines of therapy). She was treated with cyclophosphamide/etoposide and vincristine/dexamethasone/asparaginase, both of which failed. She received lymphodepletion chemotherapy which resulted in a drop, but not clearance of peripheral blasts, followed by tisagenlecleucel infusion. She experienced significant cytokine release syndrome. The patient then had progressive leukemia with peripheral blasts detected three weeks after tisagenlecleucel with lineage switch to monocytic acute myeloid leukemia (AML). Repeat genetic analysis redetected KMT2A::AFF1 fusion and NRAS (Q61R VAF 0.05, Q61K with VAF 0.40) and TP53 (R248Q with VAF 0.103, R282P with VAF 0.82) mutations (specimen time point for PDX8-2). She received subsequent AML-directed chemotherapy with cytarabine, daunomycin, and etoposide and with decitabine and venetoclax, but was unable to achieve remission and died at 14 years of age of leukemia- and coagulopathy-associated complications one year after her initial diagnosis. Patient-derived xenograft (PDX) models were created from her progressive B-ALL specimen in immunocompromised NSG mice (model ALL2184/PDX8-1) and her AML specimen in busulfan-conditioned NSGS mice (AML2311, AML2263/PDX8-2) for experimental studies as described and with informed consent in accordance with the Declaration of Helsinki via institutional review board-approved research biorepository protocols.

**Patient Vignette for PDX 5-1 (ALL1979) and 5-2 (AML2704) PDX models[1],[2]:**

ALL1979 and AML2704 PDX models: A 13 month-old male was diagnosed with CD19+ NCI HR B-acute lymphoblastic leukemia (B-ALL) with a WBC > 200,000 with 95% peripheral blasts. Cerebrospinal fluid (CSF) had macroscopic evidence of leukemia (CNS3). Cytogenetic analysis showed 46,XY with t(4;11) in 14 of 15 metaphases. Fluorescence in situ hybridization (FISH) assays detected KMT2A rearrangement in 90.5% of cells. RNA-based fusion and DNA-based NGS analyses demonstrated KMT2A::AFF1 rearrangement and IKZF1 deletion (Specimen time point for PDX5-1). The patient received four-drug induction chemotherapy on the COG

AALL1131 study and cleared his CSF, but had positive end-induction MRD at 0.035% in his bone marrow. He underwent autologous T cell apheresis after induction given high future relapse risk. He received consolidation chemotherapy as per AALL1131/VHR arm A and had negative end-consolidation MRD at 0.002%. He experienced early medullary CD19+ ALL relapse with CNS2a status at 12 months from initial ALL diagnosis. Repeat genetic analysis redetected the known KMT2A::AFF1 fusion and IKZF1 deletion and acquisition of new somatic NRAS (G12A with VAF 0.32) and TP53 (R282Sfs*25 with VAF 0.73) mutations. He received reinduction therapy on the COG AALL1331 study and had positive end-reinduction MRD at 3.2% He received several additional cycles of chemotherapy for disease stabilization, but did not achieve a stable remission. He then receive lymphodepleting chemotherapy and tisagenlecleucel infusion, following which he experienced CRS and immune effector cell associated-neurotoxicity syndrome that resolved with supportive care. He was in MRD-negative remission with CNS1 status at one month after tisagenlecleucel and had normal B cell aplasia. He subsequently relapsed at three months after tisagenlecleucel (19 months from diagnosis) with CD19-negative CD22+ B-ALL (KMT2A::AFF1, NRAS G12A 0.05, TP53 R282Sfs*25 with VAF 0.11 in the setting of low level bone marrow infiltration, and further loss-of-heterozygosity of chromosome 17). He was treated with inotuzumab monotherapy on the COG AALL1621 trial with lack of remission reinduction and persistent CD22+ B-ALL (MRD 21.6%). He underwent a second T cell apheresis, received additional chemotherapy, and was treated with CD22-directed CAR T cell (CD122CART) immunotherapy on an institutional phase 1 clinical trial at two years from diagnosis, and experienced CRS. Specimen time point for PDX 5-2 was just prior to CD122 CAR-T cell therapy. Bone marrow evaluation at one month after CD22CART demonstration lack of remission and lineage switch to monocytic AML. He received palliative azacytidine, venetoclax, and gemtuzumab[3] for life prolongation and died at three years of age of leukemia- and infection-associated complications. PDX models were created from his ALL and AML specimens for experimental studies as above.

### SUPPLEMENTAL MATERIAL AND METHODS

#### Cell lines and cultures

Human MV4;11 [KMT2A-AFF1, TP53 wild type] and derived TP53 mutant sublines were maintained in RMPI-1640 supplemented with 10% FBS and 50 U/ml P/S.
All cells were cultured in a humidified incubator at 37°C in 5% CO2.

#### MENIN/KMT2A inhibition in human AML cell lines

Cells were plated in duplicates and VTP50469 (Syndax®) or DMSO control was added at the indicated concentration. Cell growth and viability were determined by trypan blue exclusion staining after 96 hours.

#### Human samples

Samples from pediatric leukemia patients were obtained from diagnostic procedures at the Children's Hospital of Philadelphia, with patient informed consent according to the Declaration of Helsinki and institutional review board approval.

#### Human leukemia patient derived xenograft model

All experiments were conducted in accord with the principles and procedures outlined in the Internal Animal Care and Use Committee (IACUC).

NSG (NOD-scid IL2Rgnull) and NSGS (NOD-scid IL2Rgnull-3/GM/SF) were obtained from Jackson laboratories® and maintained under specific pathogen free conditions. 24-48 hours prior to transplantation, mice were conditioned with busulfan i.p. Previously banked patient leukemia samples were thawed, resuspended in PBS (Life Technologies, Carlsbad, CA, USA) and injected into the tail vein. Mice were placed on VTP-50460 or equipotent SNDX-5613 0.1% chow or control chow for the duration indicated in the respective figures. Myeloblasts were detected in peripheral blood, bone marrow and spleen after staining with a combination of anti-human CD45 (Alexa 700, BD Pharmingen, clone HI30) and anti-mouse CD45 (FITC, BD Pharmingen, clone 30-F11) antibodies.

**RNA extraction**

Leukemia cells were isolated from 3 independent moribund mice each, thawed and sorted on live (FSC vs SSC and DAPI negative), human CD45+, mouse CD45- cells. RNA was extracted using the RNeasy mini kit from QIAGEN (Hilden, Germany) according to manufacturer instructions. Briefly, gDNA was removed by a column, the flow through was mixed 1:1 with 70% ethanol, the RNA was bound to a second column and washed with ethanol. RNA was eluted in water and quantified using a NanoDrop® spectrophotometer (Thermo Fisher Scientific, Waltham, MA, USA). RNA-Seq libraries were prepared from PolyA selected mRNA (eukaryotic), and multiplexed sequencing was conducted on Illumina HiSeq Sequencers to a read depth of 20-30 million per sample.

**RNA-seq analysis**

Raw Fastq files were aligned using STAR (ref. 30) against reference Mus musculus GRCm38. All samples read-counts were quantified by Kallisto (version 0.45.0). Output from Kallisto was then directly imported into DESeq2 in order to detect differentially expressed genes (DEG). DEGs were deemed as genes with False Discovery Rate (FDR) less than 0.05. All analysis was carried using R, version 3.6.3.

Gene Set Enrichment Analysis (GSEA) was carried out using GSEA standalone software (Version 4.03) and “GSEAPreranked” tool. “Stat” field from the output of DESeq2 was used as ranked list input for GSEA software. Gene lists in 'Gene sets database' was constructed from the publications listed in ***Supplementary Table 1***. We selected "classic" as our parameter for enrichment score as suggested by the GSEA software instruction manual. RNA-Seq data has been submitted to GEO and access to the data will be provided upon request.

**Statistical analysis**

Statistical analysis of $hCD45^+$ cells in bone marrow, spleen and peripheral blood was performed by unpaired two-tailed Student’s t-tests. Statistical analysis of survival was carried out using Kaplan Meyer estimates (Prism 8 software).
